# Simulation and Sensitivity Analysis and Cross-Validation, Demonstrating the Utility of Genteract GxE Discovery methods

**DOI:** 10.1101/2020.11.25.396861

**Authors:** Brody Holohan, Raphael Laderman

## Abstract

Gene-environment interactions are at the heart of why many complex traits are not fully heritable, and why prediction of disease incidence and individual response to environmental changes based on genetics has been underwhelming in utility. Understanding these interactions is the primary limiting factor for the application of personalized medicine, but current methods are not well suited for dealing with complex traits that pose both a dimensionality and sparse data problem to unsupervised analysis methods. Genteract has developed a proprietary analytical technique that allows for detection and interpretation of GxEs regarding specific pairs of a single phenotype with a single environmental factor; these methods allow us to develop a platform that can be used to predict how individuals will respond to changes in their environment based on their genetics. To validate the methods we performed two types of testing: cross-validation against a dataset of clinical study results, and application of the methods in a simulated dataset. These tests enable a greater understanding of the methods’ utility, statistical power and predictive capabilities.

## Introduction

Individual genotype can affect the way that different individuals respond to specific environmental stimuli by altering the function, abundance or regulation of gene products influenced by a given variant in the genome. Such interactions between a given phenotype of interest, an environmental factor of interest and genotype at a specific site in the genome are collectively referred to as Gene-Environment Interactions (GxEs). GxEs are one reason why well-powered Genome-Wide Association Studies (GWASs) fail to detect variants that explain a large fraction of the genetic contribution to phenotypes such as obesity, chronic pain, insomnia, working memory and athletic ability despite the fact that twin studies intimate a large amount of genetic control of these phenotypes. The presence of many GxEs would explain this discrepancy because if many individuals responded differentially to a given environmental factor with respect to a specific phenotype based on their genetics, the direct genotype-dependence of the phenotype would be masked by the differential exposure present in the population to the environmental factor that site controls response to; GWASs studies would miss sites that control GxEs because the population as a whole has a differing response to a heterogenous stimulus, which results in a population-level random walk.

A number of methods to detect GxEs currently exist, which almost universally use the case-control logic best embodied in the Plink toolset^1,2^. Generally, the phenotype of interest is categorized into “Cases/Affecteds” and “Controls/Unaffecteds”, and a variety of statistical tests are performed to detect deviations from expected genotype frequencies between these two categories^3^. Though these methods are useful for Mendelian genetic diseases with very stark differences between affected and unaffected individuals, but are not well suited for multivariate comparisons (for instance, gene-environment-phenotype), and the logic is almost fully unable to handle interactions between continuous phenotypes, continuous environmental factors and non-allele-dose-dependent effects. Further, the binning requirements of these methods necessitate often-arbitrary distinctions between cases and controls (for instance, BMI > 25 “affected” by obesity, BMI <= 25 “unaffected by obesity”) that do not correctly reflect these phenotypes’ continuous nature. Much information is sacrificed when this inappropriate binning is conducted, and though attempts have been made to deal with this problem, for instance via permutation, these measures are computationally intensive and still introduce a number of opportunities for ascertainment and publication bias.

First, we describe the use of our method to detect GxEs in a total information context--simulated population data containing GxE interactions between a phenotype and an environment, as well as its use on a simulated population without GxEs. In order to further evaluate the accuracy, precision and reproducibility of the predictions made by our methodology, we then perform cross-validation, utilizing half the data for a clinical study dataset as a training set, and then validating the predictions created by the method on the other half of the data. This cross-validation indicates that our methodology is able to make predictions from the source data with very low false discovery rates and high real positive rates at varying levels of stringency depending on the application’s requirements.

## Simulation Methods

In this section, we discuss the use of our analysis methodology on simulated genotype, phenotype and environmental factor data to illustrate the methods’ sensitivity, specificity and a number of factors that can influence the rate of spurious results. The workflow for the simulation is shown in Figure 1.

**Figure 1:**
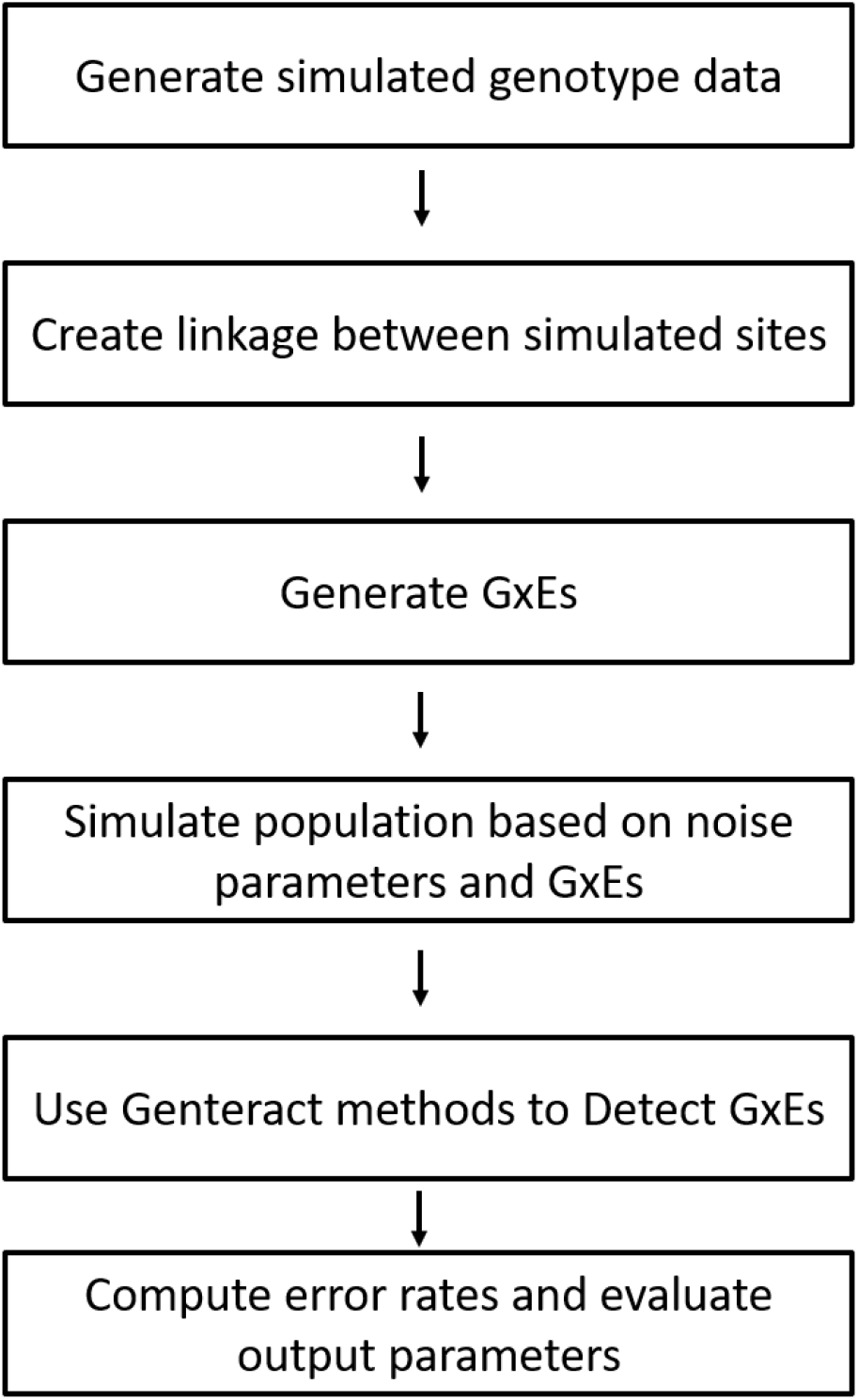
Simulated environment for sensitivity analysis of the method.

First, a simulated population of 2000 individuals is generated for 500 hypothetical sites, and linkage between these sites is enforced such that each site will conform to its specific minor allele frequency while maintaining linkage to a number of pre-specified other sites in order to simulate population stratification. See Supplemental Repository 1 and Table 1 for the code used in this process and the parameters used for this example, respectively.

**Table 1:** Parameters used in sensitivity analysis.

| numSites | proportionCausal | meanLinkedSites | sdLinkedSites | minLinkStrength | meanLinkStrength | sdLinkStrength | maxLinkStrength | randSeed | numSamp |
| --- | --- | --- | --- | --- | --- | --- | --- | --- | --- |
| 500 | 0.02 | 3 | 3 | -0.95 | 0 | 0.4 | 0.95 | 111 | 2000 |

| indMean | indSD | depMean | depSD | betaMean | betaSD | interMean | interSD | propCompliant | numComp | noiseMean | noiseSD |
| --- | --- | --- | --- | --- | --- | --- | --- | --- | --- | --- | --- |
| 40 | 40 | 100 | 1 | 0 | 0.5 | 10 | 5 | 0.9 | 45000 | 0 | 2 |

**Table 4:** Logic used to score positives and negatives for the ROC plot.

|  | Positive (call)<br>in training set | Negative (no call)<br>in training set |
| --- | --- | --- |
| Positive (responder)<br>in testing set | True positive | False negative |
| Negative (non-responder)<br>in testing set | False positive | True negative |

Next, a fraction of the simulated loci are randomly assigned “causal” for the phenotype:environment pair to be simulated, and an effect on the response to the environment in terms of the phenotype for each genotype at these sites is assigned. The simulated population’s environmental variable values are then assigned from a normal distribution with a mean and standard deviation specified in Table 1, and the population’s phenotype values are assigned from a distribution specified in Table 1 and then influenced by the total effect of their genotype at all causal loci and their exposure to the environmental variable. This generates a sample data set of genetic data that contains a small number of GxE sites and a large number of sites that are not related to the phenotype:environment pair but may be linked to GxE sites, as well as phenotype and environment data that can be used to evaluate the method’s ability to detect real GxEs and make accurate predictions about response.

Next, the methods described herein are used in the same way they are used in real datasets to detect candidate GxEs and make predictions about response. Then a simulated intervention is performed on the population, where each individual is assigned an intervention that changes their environmental variable exposure by a value sampled from a distribution specified in Table 1, and a new phenotype value is calculated based on their genotypes at GxE sites and a noise value sampled from a distribution specified in Table 1 in order to simulate a response to the differing exposure to the environmental variable as well as evaluate the method’s’ ability to tolerate noise. Each site is then scored based upon how well its beta from the method predicted each individual’s response to the environmental factor, and the method as a whole is evaluated to determine how frequently it identifies false positive and false negative results, as well as its overall ability to predict response under the assumptions evaluated.

This test can be performed in batches with different parameters in order to explore the method’s utility in different regions of the parameter space, as well as to test how modifications to the methods modify their ability to make accurate predictions.

## Simulation Results

For this simulation, 500 sites were considered, of which 10 (2%) were flagged as causal. Each site had on average 3 enforced links to other sites, with a standard deviation of 3 links per site. 2000 simulated individuals were created with genotypes at each of the 500 sites, and linkage between these sites was created through a weighted probability of minor allele method enumerated in greater detail in supplemental repository 1. The initial mean of the independent variable was 40, with a standard deviation of 40, and the initial mean of the dependent variable was 100 with a standard deviation of 1. The GxEs at causal loci had a mean effect size of 0 with a standard deviation of 0.5, and the intervention to be evaluated had a mean value of 10 units of the independent variable with a standard deviation of 5 units. For this simulated intervention, 90% of simulated subjects were compliant, and 45000 inter-individual comparisons were conducted in the GxE discovery setting. Noise with a mean of 0 and a standard deviation of 2 was applied after the intervention to represent the effect of other variables as well as other sources of randomness.

Using the GxE discovery analysis described above, the method made a correct prediction about the direction of response 1375 times out of 1828 individuals who were compliant with the simulated intervention (75.21%). It is notable that this figure includes individuals with very small predicted effects, which led to the development of the thresholding method used below in the cross-validation; this implements a region wherein no prediction is made about an individual because of the weak statistical significance of the trend.

In the simulation 3 of the 25 sites that were significant after correcting for the number of tests performed (p <= 1*10^-4, Figure 2, top left)) were causal, and all three significant causal sites outperformed the significant non-causal sites in the intervention (made more correct predictions). All three of the causal sites were starkly demarcated from non-causal sites following the intervention, as they made correct predictions a much larger proportion of the time than the non-causal sites (Figure 2, top right, causal sites are triangular). 7 of the causal sites were not significant in the initial discovery analysis, but two of these would have been readily apparent following the intervention, demonstrating that it is possible to use the data in an intervention to recover false negative results of the initial GxE discovery. A linear regression of the change in dependent variable on change in environmental variable multiple by genotype score (the prediction of response. Figure 2, bottom) generated an r^2^value of 0.5487, indicating that the method had substantial predictive power even before the response data was used to improve the predictions.

**Figure 2:**
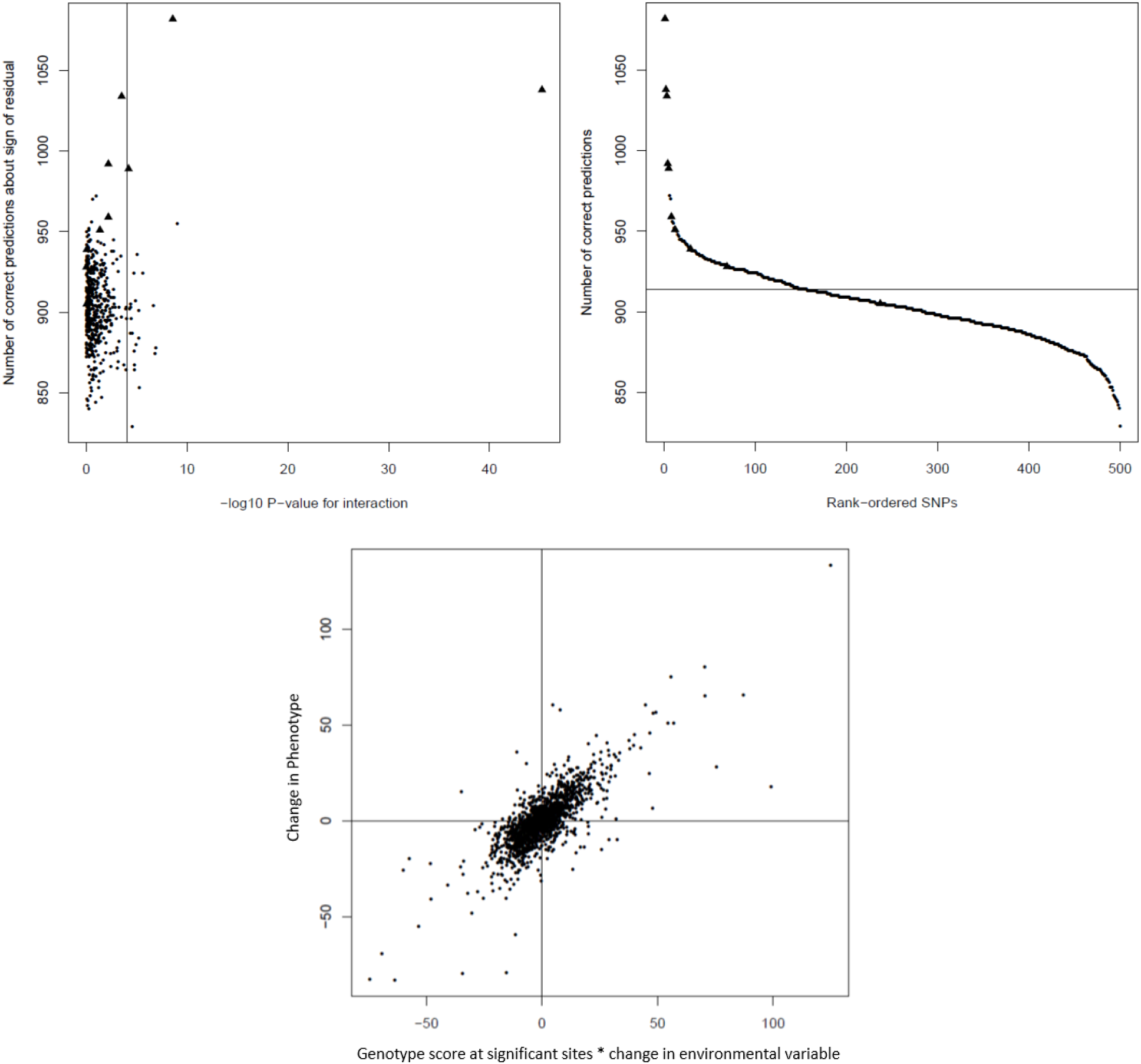
Representative results of use of the method in simulated data.

To determine the rate of false positives in the presence of no causal loci, the simulation and analysis procedure described above was repeated with the same parameters, except that the proportion of simulated loci assigned to be causal was 0; any changes in phenotype in the simulated intervention were due to noise alone, and there was no relationship between the phenotype and the environmental factor. No loci in this analysis were significant above the cutoff accounting for the number of loci considered (1*10^-4, Figure 3, top left). The number of correct predictions made by all sites was centered around the rate expected by chance (Figure 3, top right), and no site made correct predictions at the same rate as the top causal loci in the iteration of the simulation that did have causal loci. Further, predictions made by the analysis considering all loci (Figure 3, bottom center) were unrelated to the actual response.

**Figure 3:**
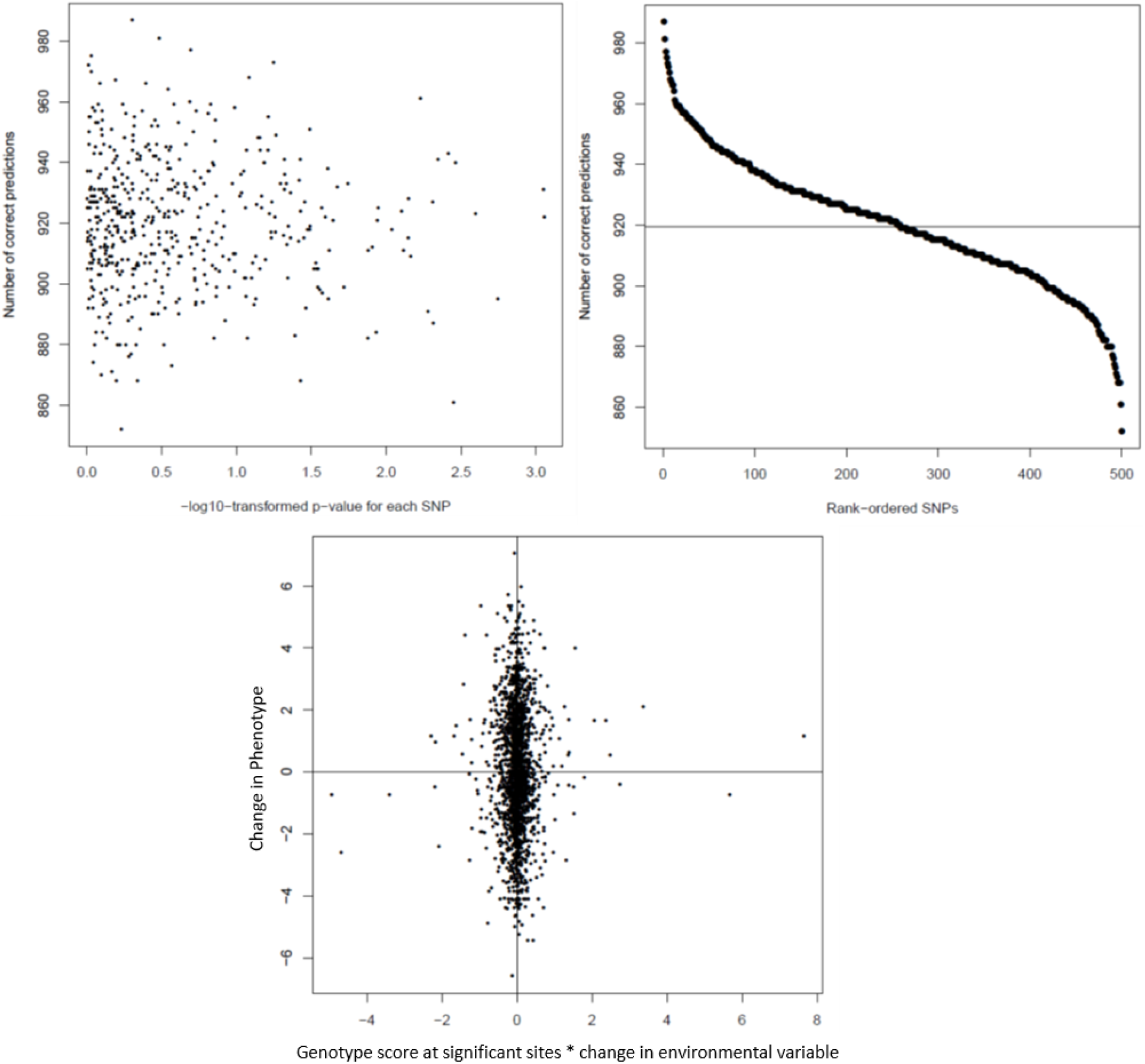
Use of the method on simulated data with no causal loci.

## Cross-Validation Methods

In order to evaluate the accuracy, precision and reproducibility of the predictions made by our methodology, we perform cross-validation, utilizing half the data for a clinical study dataset as a training set, and then validating the predictions created by the method on the other half of the data. The overall procedure followed is detailed in Figure 4, below. This cross-validation indicates that our methodology is able to make predictions from the source data with very low false discovery rates and high real positive rates at varying levels of stringency depending on the application’s requirements.

**Figure 4:**
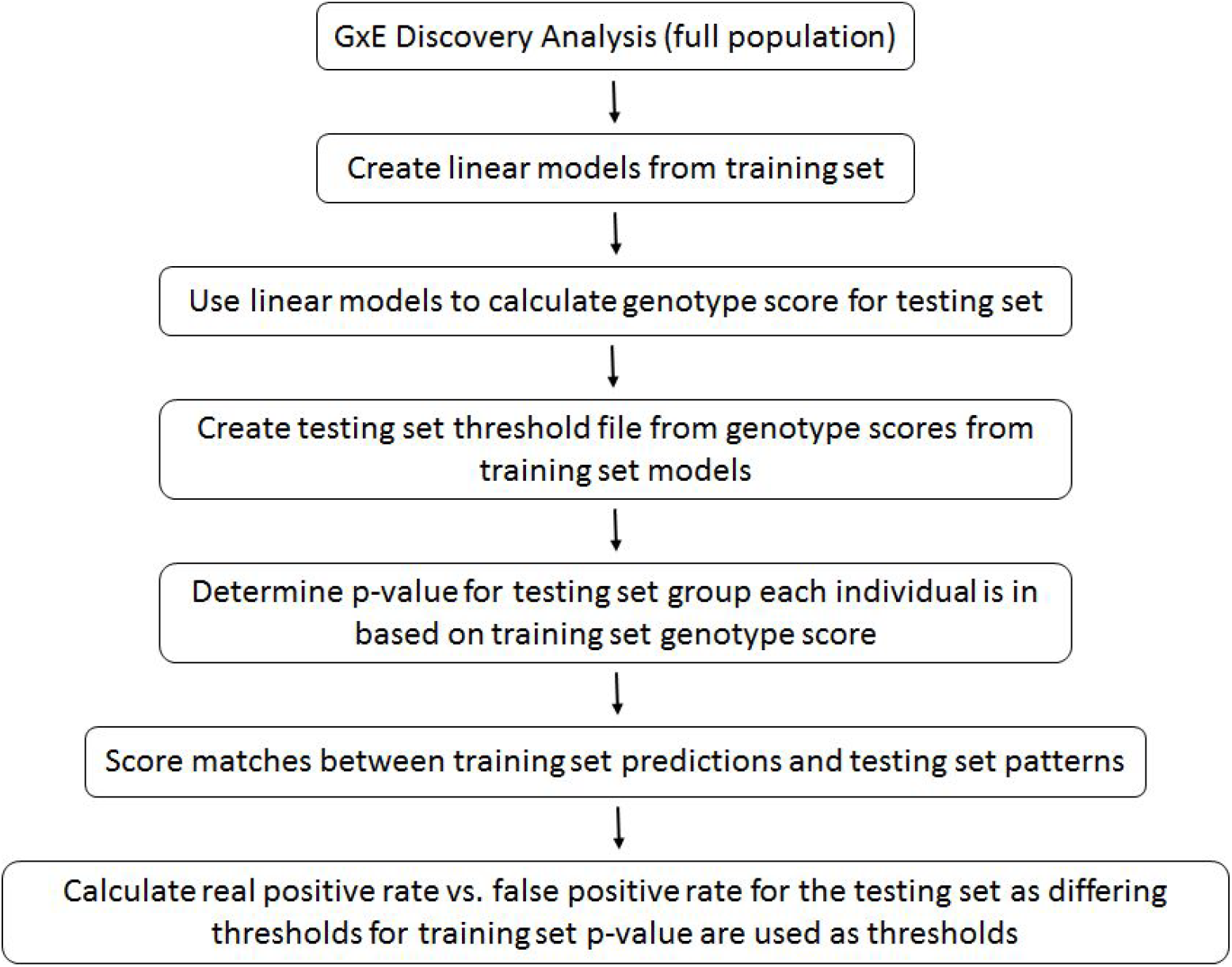
Procedure used in cross-validation.

Genteract GxE discovery methods were used to identify GxE relevant sites for a Phenotype:Environment (P:E) pair of interest (BMI : Dietary Maltose intake) using data from the entire population of the Multi-Ethnic Study of Atherosclerosis (MESA). Next, the population was split into a training set and a testing set at random with a 50:50 ratio, and linear models were constructed for each genotype at each locus of interest using BMI as the dependent variable and Maltose Intake as the independent variable for all possible genotypes at each site of interest. These linear models were used to create a “strength file” consisting of the beta value for each possible genotype at each locus of interest, which was then used to calculate a “genotype score” for each individual in the training set using Genteract methods.

The training set (1,933 individuals) was sorted in ascending order by genotype score. 22 population sections were then created sorted by genotype score, each section consisting of a moving window 300-individuals, with the window moving up in genotype score by 75 individuals with each increment. Moving linear models of BMI vs. Maltose intake for each were then constructed for the each section in the training set, and the resulting summary data was stored in a threshold file, which summarizes a linear model of the dependent variable (BMI) on the independent variable (maltose intake) in the subset of the population evaluated for each segment. Next, the strength file from the training set was used to calculate a genotype score for each individual in the testing set, and the same procedure was used to create a threshold file for the testing set.

In order to evaluate the specificity and sensitivity of the methods and use classifier evaluation logic to inform on the utility of the methods, it is necessary to decide what is a correct or incorrect prediction. In this case, if an individual in the testing set was in a section of the moving linear model with a significant trend (p < 0.01), they were scored as a positive, indicating that that individuals segregated by genotype score calculated from the training set had a meaningful relationship between the phenotype and the environment. Individuals in the testing set in a section of the moving linear model without a significant trend were scored as negatives.

The moving linear models from the training set were used to evaluate if the methods made a call about an individual in the testing set by determining which section of the moving linear model in the training set had a genotype score closest to a given individual in the testing set; the classifier evaluated is the p-value of the moving linear model section in the training set, and the threshold for this p-value to make a call.

As such, a true positive was scored when an individual in the testing set with a significant p-value from the training set threshold file (a call) also had a significant p-value from the testing set threshold file (a positive). A false positive was scored when an individual in the testing set with a significant p-value from the training set threshold file did not have a significant p-value from the testing set threshold file (a negative).

A true negative was scored when an individual without a significant p-value from the training set (no call) also did not have a significant p-value from the testing set threshold file, and a false negative was scored when an individual without a significant p-value from the training set did have a significant p-value from the testing set threshold file (Table 3).

## Cross-Validation Results

Varying the cutoff used as a classifier from p < 1 to p < 1*10^−10^ resulted in the ROC plot shown in Figure 5. At all ranges the real positive rate was higher than or equal to the false positive rate, and very favorable real positive rate:false positive rate ratios were achieved when the classifier cutoff was set at p< 1*10^−6^. The cutoff can also be adjusted to completely eliminate false positives at the expense of a reduced overall prediction rate.

**Figure 5:**
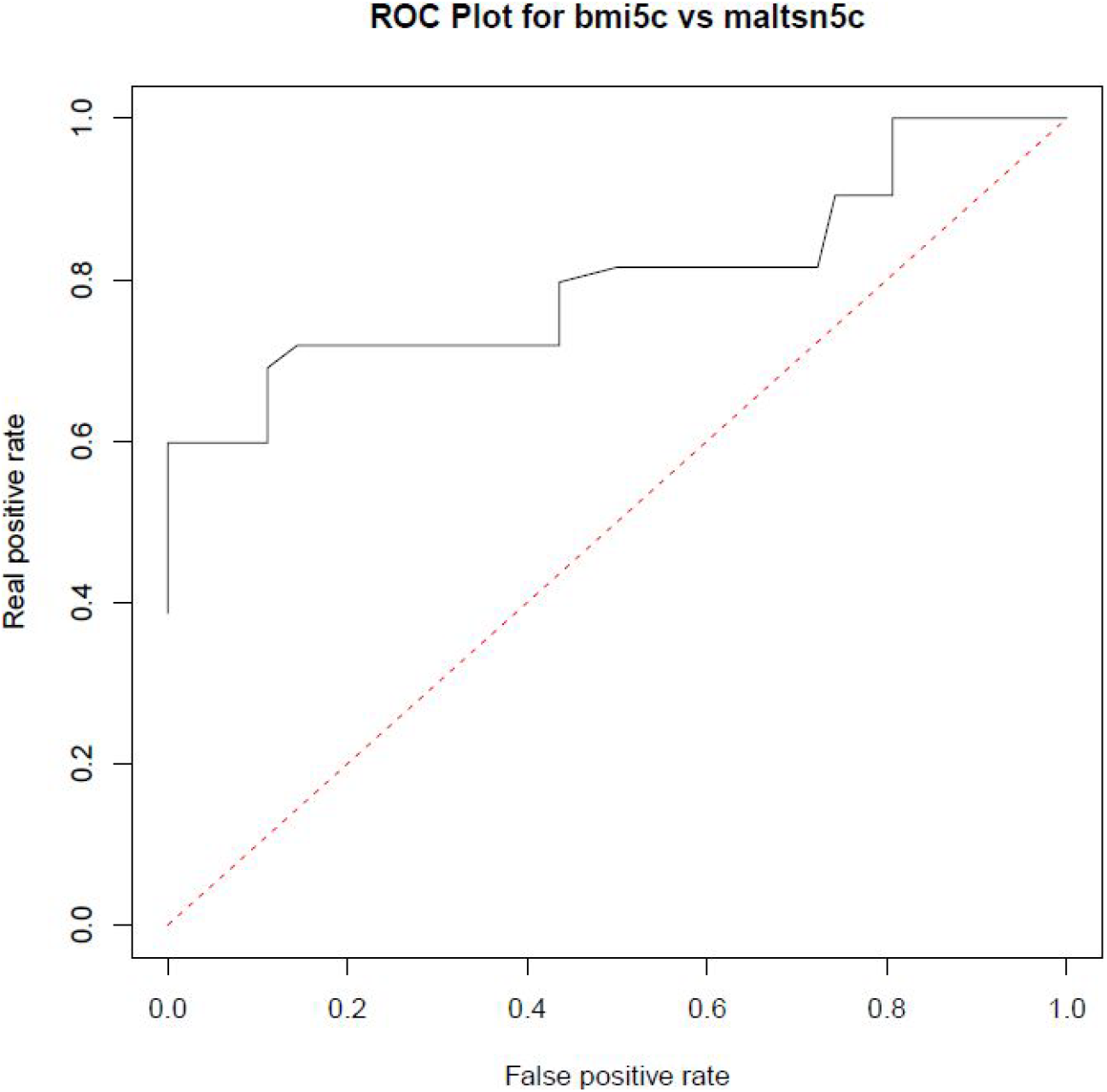
ROC plot for BMI vs. Maltose intake in the MESA Cohort.

## Discussion

Genteract’s proprietary gene-environment interaction analysis methods can detect causal loci that govern gene-environment interactions in both simulated data as well as in real data from large studies in human populations. We have utilized these methods to tune the logic used to make predictions about an individual’s response to environmental factors based on their genetics and retrospectively validated this logic in large clinical trial data.

The simulated use of these methods is provided in order to demonstrate that they are capable of detecting and correctly interpreting gene-environment interactions when they are present and the correct conditions are present (noise, relative strength of the associations, sample size) in the context of a system where objective ‘ground truth’ correct answers are detectable.

Further, these procedures are used to set priorities for the prediction algorithms; for instance, where sensitivity is preferred over specificity, a classifier cutoff that is more tolerant of false positive results can be selected in order to optimize the real positive:false positive ratio (mathematically represented by minimum distance from the cartesian point 0,1 on the ROC plot). For applications that prefer specificity over sensitivity, for instance in the prediction of a favorable response to a drug, a classifier cutoff that minimizes the rate of false positives first and then optimizes for real positives can be used.

The methods described here have been validated on and are initially being utilized for the discovery and validation of gene-environment interactions regarding low-risk environmental factors such as foods, nutritional supplements and noninvasive medical devices. The same methods are also applicable to prescription and over-the-counter medicines for which source data is available, and can be applied to guide the development of experimental medicines when utilized in tandem with a traditional clinical trial.

## References

1. Purcell, S., B. Neale, K. Todd-Brown, MAR Ferreira, D. Bender, J. Maller, P. Sklar, PIW De Bakker, MJ Daly, and PC Sham. 2007. “PLINK: a toolset for whole-genome association and population-based linkage analysis.” American Journal of Human Genetics 81. https://pubmed.ncbi.nlm.nih.gov/17701901/.

2. Purcell, Shaun. “PLINK.” http://pngu.mgh.harvard.edu/purcell/plink/.

3. Kaler, Avjinder S., and Larry C. Purcell. 2019. “Estimation of a significance threshold for genome-wide association studies.” BMC Genomics 20. https://doi.org/10.1186/s12864-019-5992-7.

